# Building a barrier: investigating the functional role and evolution of induced trichome production

**DOI:** 10.1101/2023.11.16.567395

**Authors:** Nia M Johnson, Regina S Baucom

## Abstract

Functional traits are increasingly recognized as fundamental determinants for mediating ecological resilience. As global change accelerates, phenotypic plasticity in functional traits may provide a modality for fine-tuning responses to atypical or stressful environments. While various studies have explored induced plant trichomes as a response to biotic stress, emerging evidence suggests that trichome induction may also function as a broader tactic for managing a range of environmental stressors, including abiotic chemical stress. Using the annual invasive weed velvetleaf (*Abutilon theophrasti*), we examine whether herbicide exposure induces changes in trichome traits and whether these responses are associated with herbicide resistance and fitness outcomes. We quantified changes in trichome density and trichome type proportions across control and glyphosate (active ingredient in “Roundup”) treated plants. We identified positive correlations between induced total trichome density and herbicide resistance, as well as induced branched trichomes and herbicide resistance. Selection analysis further revealed positive linear selection acting upon induced trichome density in the presence of herbicide, as well as correlative selection favoring induced trichome density and intermediate plant growth. Overall, our study indicates that plastic defenses like trichomes may contribute to plant performance under novel abiotic stress and highlights key constraints shaping the evolution of stress-responsiveness.

## Introduction

Understanding the relationship between functional traits and environmental shifts is a growing area of both evolutionary and ecological research given that global change is intensifying the frequency and severity of stressors (Aubin et al 2016, Ahrens et al 2019, Madsen-Hepp et al 2023). Phenotypic plasticity, the ability of a genotype to express various phenotypes as a response to environmental heterogeneity (Sultan and Stearns 2005), is a proposed mechanism by which plant species may modulate functional traits and persist amidst rapid environmental change (Valladares et al 2007). As abiotic challenges such as temperature extremes, altered precipitation regimes, and chemical exposures escalate, investigating how plastic functional trait responses are deployed across environments can reveal important insights into processes influencing population resilience.

Among functional traits, plant trichomes—epidermal outgrowths of various types—are widely recognized for their protective roles in responding to both biotic and abiotic stress. Although often studied as a defense against herbivory (Doss et al 1987, Haberlandt 1914, Agren and Schemske 1993, Westerbergh and Nyberg 1995, Dalin and Bjorkman 2003), trichomes also respond to abiotic factors such as ultraviolet radiation (Tattini et al 2000, Yamasaki et al 2007), drought (Boughalleb and Hajlaoui 2011), elevated CO_2_ (Karowe and Gribb 2011), and temperature (Liakopoulos et al., 2006). These responses suggest that trichome production may be a general stress-responsive trait, adjusting to diverse environmental cues. However, the degree to which different trichome types respond plastically to novel stressors, such as herbicide exposure, is still poorly understood.

As chemical agents applied across agricultural (Gianessi 2013), urban (Meftaul et al 2020), and restoration landscapes (Humphries et al 2020), herbicides represent a particularly relevant and underexplored abiotic stressor. There is sufficient evidence that herbicide drifts into surrounding ecosystems (Kleijn and Snoeijing 1997, Marrs et al 1989, De Snoo and Van der Poll 1999, Egan and Mortensen 2012, Hwang et al 2022), unintentionally exposing non-target plants to chemical stress (Egan & Mortensen, 2012, Silva et al 2016). Recent uptakes in herbicide use (Benbrook 2016) also highlight the crucial need to understand its impacts on population and community dynamics. Though some studies have examined herbicide effects on plant growth and herbivory (Dewar et al., 2000; Wu et al., 2001), the extent to which herbicide exposure induces plastic changes in plant defense traits such as trichomes—and whether these traits play a role in herbicide resistance and plant fitness—remains unclear.

Here, we explore the extent to which trichome traits respond plastically to herbicide exposure and examine the potential functional significance and evolutionary trajectory of such responses. Using the model weed species *Abutilon theophrasti* (hereafter velvetleaf), a species with diverse trichome types and a history of herbicide exposure in agricultural settings, we assess whether trichome traits are induced under herbicide stress, and whether such plastic changes correlate with resistance and/or fitness outcomes. There are four types of trichomes on the surface of velvetleaf, including two multicellular types that are glandular (peltate and capitate) and two unicellular types that are non-glandular (single and branched) (Sterling and Putnam 1987). While we know that glandular trichomes have the ability to synthesize and secrete flavonoids as chemical defense (Sterling et al 1987), limited work has explored the defensive role for the non-glandular types in this species. Further, velvetleaf has evolved reduced susceptibility to herbicides routinely used in weed management, including atrazine (Anderson and Gronwald 1991) and glyphosate (Hartzler and Battles 2001), but the mechanisms underlying this reduced susceptibility are unknown. Thus, for both of these reasons – *i.e.*, variation in trichome types and reduced herbicide susceptibility – velvetleaf provides a unique study system with which to address questions about stress-induced functional trait responses.

In past work, we found positive correlations between constitutively produced branched trichomes and glyphosate resistance, indicating that this trichome type may serve as an herbicide resistance trait in velvetleaf (Johnson and Baucom 2024). We further showed positive selection on branched trichomes in the presence of the herbicide, indicating that herbicide exposure could lead to adaptive evolution of this trichome type. We also observed that trichome phenotypes exhibited high plasticity between growth room and field conditions, and reasoned that, similar to trichome induction when plants are damaged by herbivory or abiotic factors, damage from herbicide may induce trichomes in surviving plants. In this work, we investigate trichome plasticity in *Abutilon theophrasti* and examine potential impacts of trichome induction on plant fitness and performance.

We address the following specific questions: Does the density or proportion of different trichome types (branched, single, capitate, peltate) vary when exposed to herbicide in field conditions, such that trichome traits are induced in *Abutilon theophrasti* exposed to herbicide? Is there a relationship between induced trichome traits and herbicide resistance, suggesting that trichome induction may be a general feature of more resistant plants? Further, we wanted to assess the evolutionary trajectory of induced trichome defense. Thus, we asked: Is there genetic variation associated with induced trichome traits, and do plants that display trichome plasticity also exhibit higher fitness in the presence of herbicide, indicating a benefit of trichome induction? Are there fitness costs associated with trichome plasticity, and/or tradeoffs between this form of defense and growth or reproduction? The presence of genetic variation in induced trichome traits would provide evidence that induced trichome traits have the potential to evolve, whereas a benefit or cost of trichome plasticity would provide evidence for increases or brakes on this form of plant defense.

## Materials and Methods

### Study organism

Velvetleaf is an invasive annual native to Asia and is frequently found in and around corn, cotton, and soybean fields globally (Warwick and Black 1985). First introduced to Pennsylvania and Virginia in the late 17th century, velvetleaf was originally cultivated as a commercial fiber crop (Spencer 1984). Despite its commercial success in other regions, the fiber industry in the US was not able to use velvetleaf effectively, leading to the lack of management and rapid growth in and around agricultural fields. Today, this annual species has become one of the most detrimental weed species between 32° and 45° N latitude, particularly in southwestern Canada and the United States (Warwick and Black 1985).

Velvetleaf is a highly self-pollinating, hexaploid species with 2n = 42 chromosomes, and with high levels of fixed heterozygosity as a result of polyploidy (Warwick and Black 1985). Seed production ranges from 700 to 17,000 seeds per individual (Winter 1960; Chandler and Dale 1974; Hartgerink and Bazzaz 1984; Warwick and Black 1985), and seed dormancy can last up to 50 years (Warwick and Black 1988). Studies on velvetleaf accessions in North America reveal phenotypic variation that correlates with latitudinal environmental heterogeneity. For instance, southern populations found in North America produced fewer, larger seeds with lower rates of seed dormancy when compared to northern populations (Warwick and Black 1985). This species has also been shown to exhibit resistance or decreased susceptibility to several herbicide classes including triazine (Ritter 1986, Anderson and Gronwald 1991) and glyphosate (Hartzler and Battles 2001).

In addition to resistance to herbicides, velvetleaf also has traits that reduce the potential for herbivory. These traits include glandular trichomes on the plant surface, which serve as chemical defense structures via the synthesis and storage of flavonoids such as anthocyanin, quercetin, kaempferol, and myricetin (Morris and Wang 2013). The glandular morphs are multicellular structures that include peltate trichomes, which contain 4 - 5 cells, and capitate trichomes, which contain 12 - 15 cells (Sterling 1987, Fig 1). Velvetleaf also has non-glandular trichomes, which serve as physical defense structures by shielding the plant body. The non-glandular morphs are unicellular structures that include single trichomes, which grow perpendicular to the plant surface, and branched trichomes, which grow 4 - 8 arms (Sterling 1987, Fig 1).

**Figure 1.**
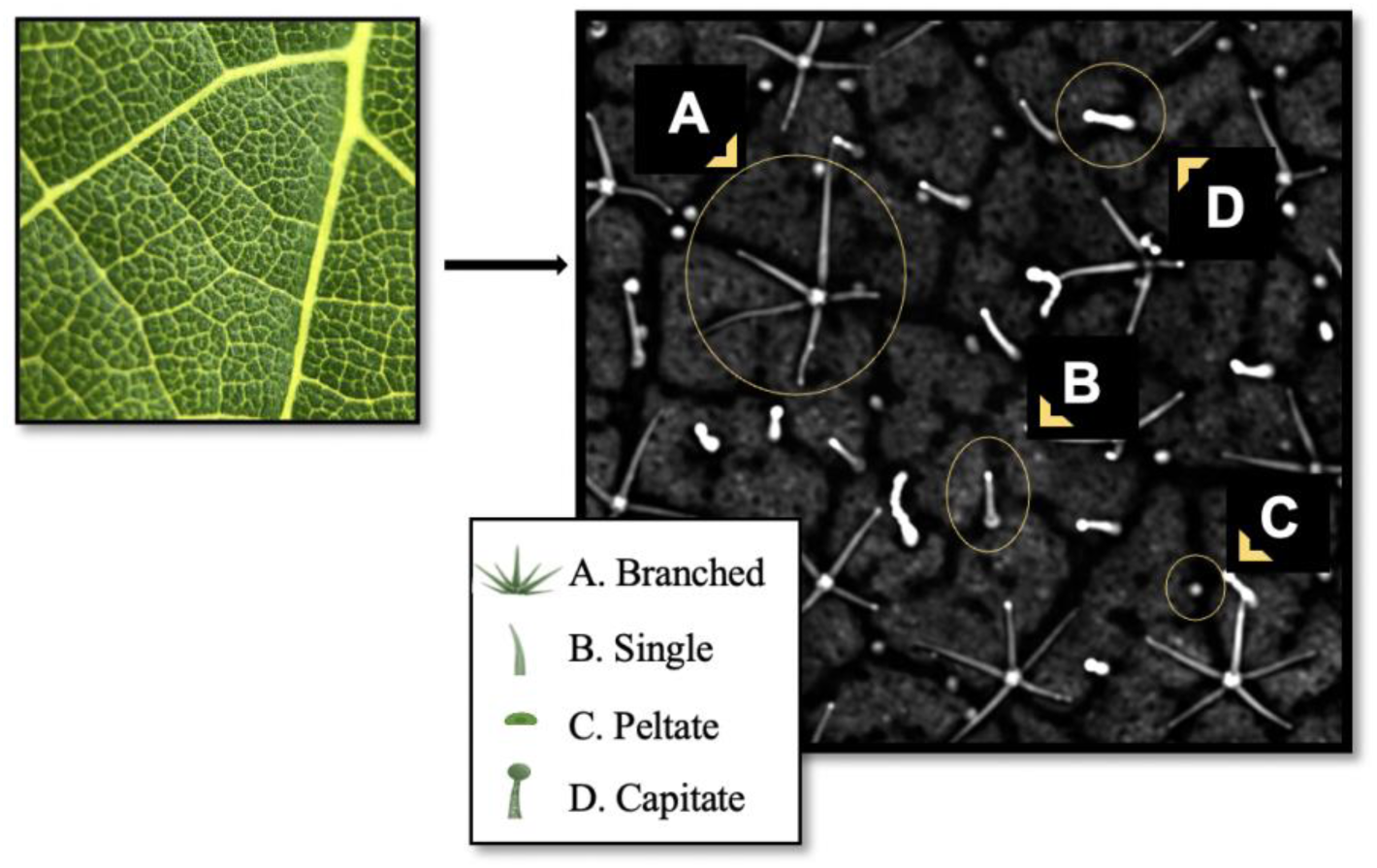
Trichome structures present on the adaxial surface of *Abutilon theophrasti* leaves using confocal imaging (x10). A) Branched trichomes are unicellular structures with 4-8 appendages. B) Single trichomes are unicellular unbranched structures. C) Peltate trichomes are multicellular glandular structures made up of 4-5 cells. D) Capitate trichomes are multicellular glandular structures made up of 12-15 cells. Density is calculated as the total number of all trichomes captured in the image.

### Field Experiment

We conducted a field experiment to measure induced trichomes in field conditions and in response to herbicide application. Seeds were sampled from individuals found in populations located near agricultural farms in Dexter, MI (see SFig 1 of Johnson and Baucom, 2024) and were planted and selfed in growth room conditions to generate replicate, selfed seed for the field experiment (Johnson and Baucom 2024). We planted scarified velvetleaf seeds in a randomized block design comprising two treatments (herbicide and control). Three replicate individuals per maternal line were planted in each treatment/block combination for a total of 720 seeds (40 maternal lines x 2 treatments x 3 blocks x 3 replicates). When plants reached an average height of 11 cm, roughly 5 weeks after germination, we applied glyphosate at 550g/ha ai to the herbicide treatments, as our previous study revealed genetic variation for reduced susceptibility at this application dose (Johnson and Baucom 2024). We recorded height measurements to estimate growth rate two weeks after planting and once again when we collected fitness estimates, approximately 12 weeks after planting.

As an annual species, velvetleaf seed count indicates the success for which individual genetic material may be passed on to subsequent generations, thus we elected to use seed count as an estimate of fitness. Plants that germinated but did not survive herbicide exposure were given a fitness of zero. To estimate herbicide damage in the field, two weeks after herbicide application we recorded the number of yellowing leaves and divided that by the total number of leaves per individual. Because velvetleaf begins to shed its leaves during seed maturation, we choose this metric to estimate herbicide damage rather than biomass so that fitness data could also be collected. Herbicide resistance was defined operationally as the inverse of herbicide damage (1 minus the proportion of yellowing leaves).

Due to low germination rates and mortality following herbicide application in the field, there were 25 maternal lines remaining for which induced trichomes could be estimated (i.e. represented in both treatments). Once plants began to flower, approximately 10 weeks after germination, we collected one leaf from 2-3 randomly chosen plants per maternal line in each treatment for trichome imaging (n = 130), ensuring that we sampled leaves that were newly formed post-herbicide exposure. Along with measuring total trichome density, we calculated the proportion of each trichome type by dividing the absolute number of each trichome structure by the total trichome density recorded in each 3mm^2^ segment. We estimated the weekly growth rate as the difference in height measured 2 weeks after germination and 12 weeks after germination divided by 10.

### Data Analysis – Phenotypic plasticity in trichome production

We conducted all statistical analyses in R (version 3.4.2, R Development Core Team). We ran a series of univariate models to assess the potential for plasticity in trichome production in different environments. For each model, the response variable was one of the observed trichome traits (density and the proportion of each type) and the fixed effect was the environment (*i.e.* control vs herbicide). We performed subsequent post-hoc tests using Tukey to detect differences between environments.

### Genetic variation underlying induced trichome traits

We tested for genetic variation underlying induced trichome traits to determine if trichome induction would be expected to respond to selection and evolve. To do so, we fit the following model for each trait:

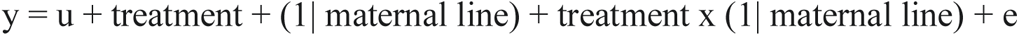

where each respective trichome traits (total density, and the proportion of branched, single, peltate, and capitate) was the response variable, u is the intercept or mean of the model, treatment was a fixed effect, maternal line was a random effect, maternal line by treatment interaction was a random effect, and e is the error term. Our specific effect of interest was the interaction of maternal line with treatment, since such an interaction would indicate that trichome traits of maternal lines responded differently to the control and herbicide environments and thus implicate adaptive plasticity. As such, the remaining analysis was performed on trichome traits with significant maternal by treatment effects. For each trait, we first removed the effects of block by performing a regression of the trait with the block effect as the sole predictor variable, and then used the residuals in each subsequent analysis of variance. We determined the significance of predictor variables using the stats package (R Core Team 2022) to generate *F*-statistics for the fixed effects. We used the lmerTest package (Kuznetsova et al 2017) to perform a log-likelihood ratio test for each random effect.

### Herbicide resistance and induced trichomes

We calculated induced trichomes in field settings as the mean difference between damaged (*i.e.* herbicide treatment) and control (*i.e.* control treatment) states for each maternal line in the field experiment. As such, this measurement produced a positive number when trichomes increased (*i.e.*, induced resistance) in response to herbicide and a negative number when trichomes decreased (*i.e.*, induced susceptibility).

We elected to perform the remaining analysis on induced total density, and induced changes in the proportion branched and capitate trichomes because these three traits exhibited significant genetic variation (see Results). We estimated Pearson correlation coefficients to determine if induction of these traits had a significant relationship with herbicide resistance and tested the significance of the correlation coefficient using the stats package.

### Genotypic selection for induced trichomes

We performed genotypic selection analysis to determine if trichome induction in response to herbicide exposure is under selection and to assess if there was a cost when compared to control conditions. We estimated differentials to measure selection on induced trichome density, proportion branched and capitate trichomes in the presence and absence of herbicide. Because the proportion of branched trichomes and capitate trichomes are not independent of total trichome density, we estimated selection differentials rather than gradients. We estimated selection differentials (*S*) by performing univariate regressions of relative fitness in each trait separately which captures total (indirect and direct) selection acting upon each trait. Relative fitness was calculated as the final seed count per individual divided by mean seed count for each treatment.

Inducible defenses are hypothesized to come at the expense of plant growth as a form of cost (Herms and Mattson 1992), thus, we explored if there was correlative selection acting on the interaction between growth rate and induced trichomes. We performed non-linear selection analyses (*γ*) containing trichome traits and growth rate as linear terms, quadratic terms, and the cross-product of the focal trichome trait and growth rate. We assessed the potential of quadratic or correlative selection by doubling the quadratic regression coefficients. Significant quadratic terms would indicate non-linear (disruptive or stabilizing) selection acting upon growth rate or induced trichome traits, whereas significant correlative selection would indicate selection favoring a combination of the focal traits.

## Results

Trichome production in *Abutilon theophrasti* is highly plastic, with significant changes in trichome density produced according to environment. When comparing between the control (non-herbicide) environment and herbicide environment in the field, plants exposed to herbicide generally exhibited fewer total trichomes (*i.e.* reduced trichome density: Fig 2A). We found no evidence for changes in the proportions of the four trichome trichome types between control and herbicide environments (Fig 2B-E).

**Figure 2.**
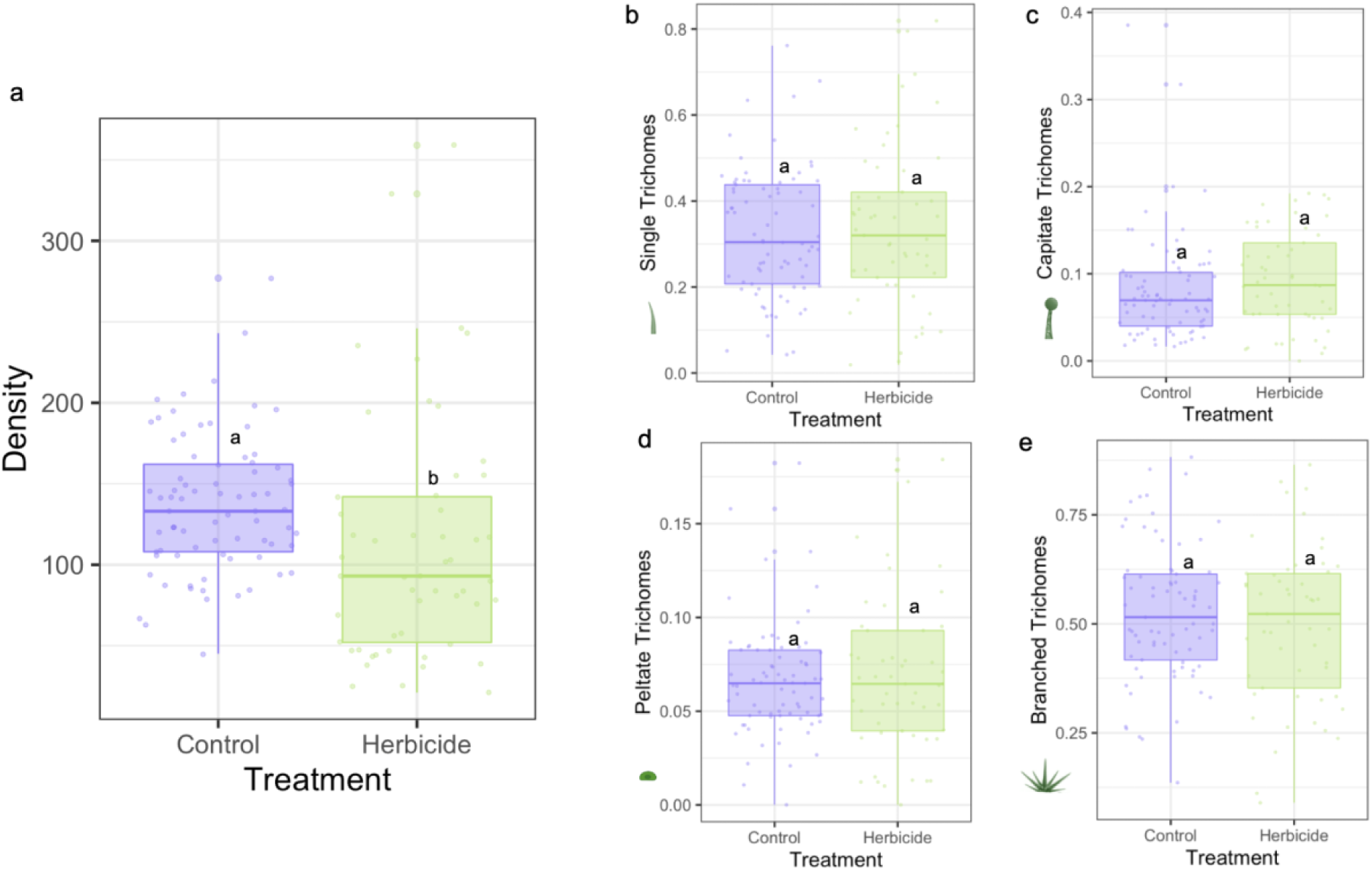
Treatment effects observed on (**a**) trichome density (F = 15.81, p = 0.02) (**b**) the proportion of single trichomes (F = 0.21, p = 0.65), (**c**) the proportion of capitate trichomes (F = 1.11, p = 0.29), (**d**) the proportion of peltate trichomes (F = 0.20, p = 0.65), and (**e**) the proportion of branched trichomes (F = 0.87, p = 0.35) in *Abutilon theophrasti*.

However, induced trichome density varied by maternal line when plants were exposed to herbicide (STable 1). We identified genetic variation in induction of the proportion of branched trichomes (maternal line by treatment effect: χ^2^ = 3.92, p = 0.048, Table 1, Fig 3), and the proportion of capitate trichomes (maternal line by treatment effect: χ^2^ = 5.64, p = 0.018, Table 1, Fig 3). Of the 25 total maternal lines, 9 increased the proportion of branched trichomes (36%), 19 increased the proportion of capitate trichomes (76%), and 6 increased the total trichome density (24%) after herbicide treatment.We also found significant maternal line by treatment interaction for total trichome density (χ^2^ = 10.05, p = 0.002, Table 1, Fig 3).

**Figure 3.**
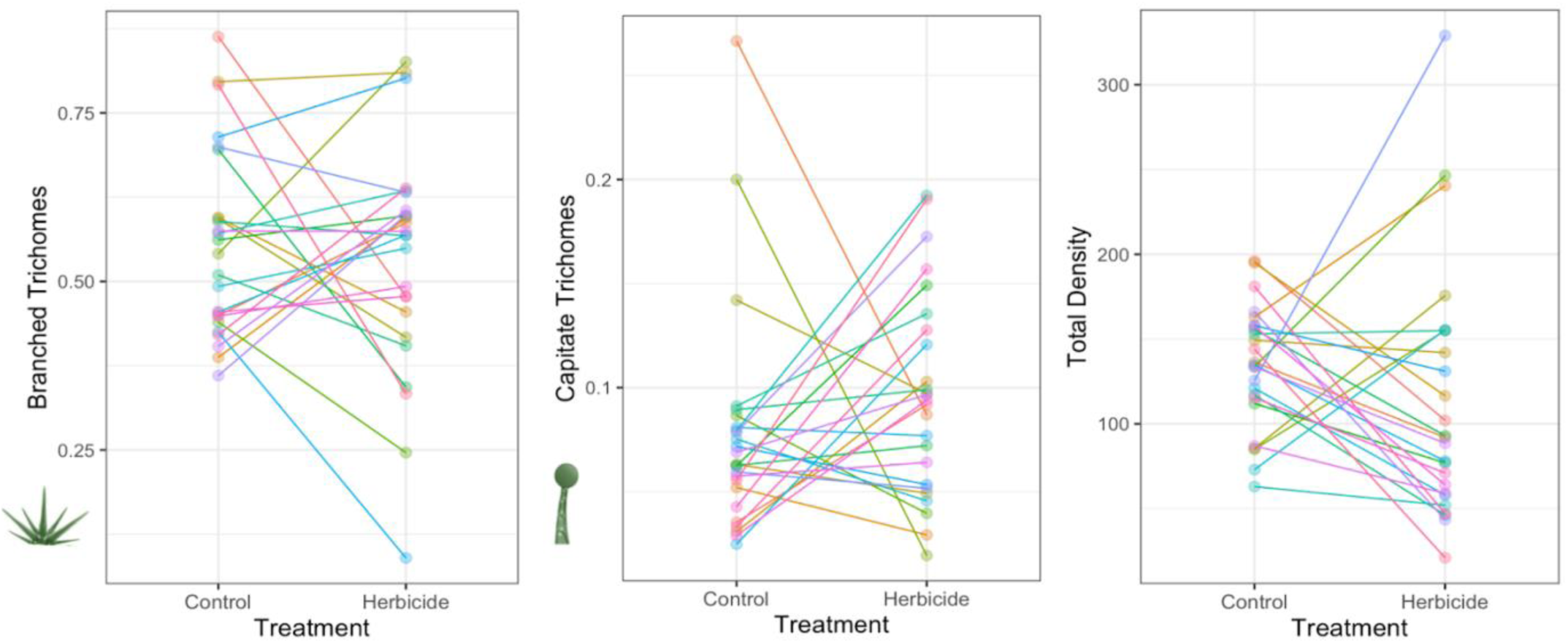
Illustration of genetic variation for induced trichome phenotypes in *Abutilon theophrasti* maternal lines grown in field conditions in either the control (non-herbicide environment) or herbicide present environment. Shown are the significant maternal line by treatment effects for the proportion branched trichomes (χ^2^ = 3.92, p = 0.048), the proportion capitate trichomes (χ^2^ = 5.64, p = 0.018), and total trichome density (χ^2^ = 10.05, p = 0.002).

**Table 1.** Results from the test for genetic variation in *Abutilon theophrasti* showing the effects of maternal line by treatment on trichome traits (proportion branched, single, capitate, peltate, and total density) captured in the field. Significant effects are indicated in boldface. The full ANOVAs are presented in STable 1.

|  | Maternal Line x Treatment Effects |  |  |
| --- | --- | --- | --- |
| | $\chi^2$ | df | p - values |
| Proportion Branched | **3.92** | **1** | **0.048** |
| Proportion Single | 3.26 | 1 | 0.071 |
| Proportion Capitate | **5.64** | **1** | **0.018** |
| Proportion Peltate | 0.71 | 1 | 0.399 |
| Total Density | **10.05** | **1** | **0.002** |

### Herbicide resistance and induced trichomes

We investigated if there was a significant relationship between induced trichomes and herbicide resistance (1 – proportion herbicide damage). We found that lines that exhibited a greater induction of branched trichomes and a greater induction of total trichome density exhibited higher levels of herbicide resistance (r = 0.44, p = 0.035, Fig 4A; r = 0.43, p = 0.040, Fig 4C).

**Figure 4.**
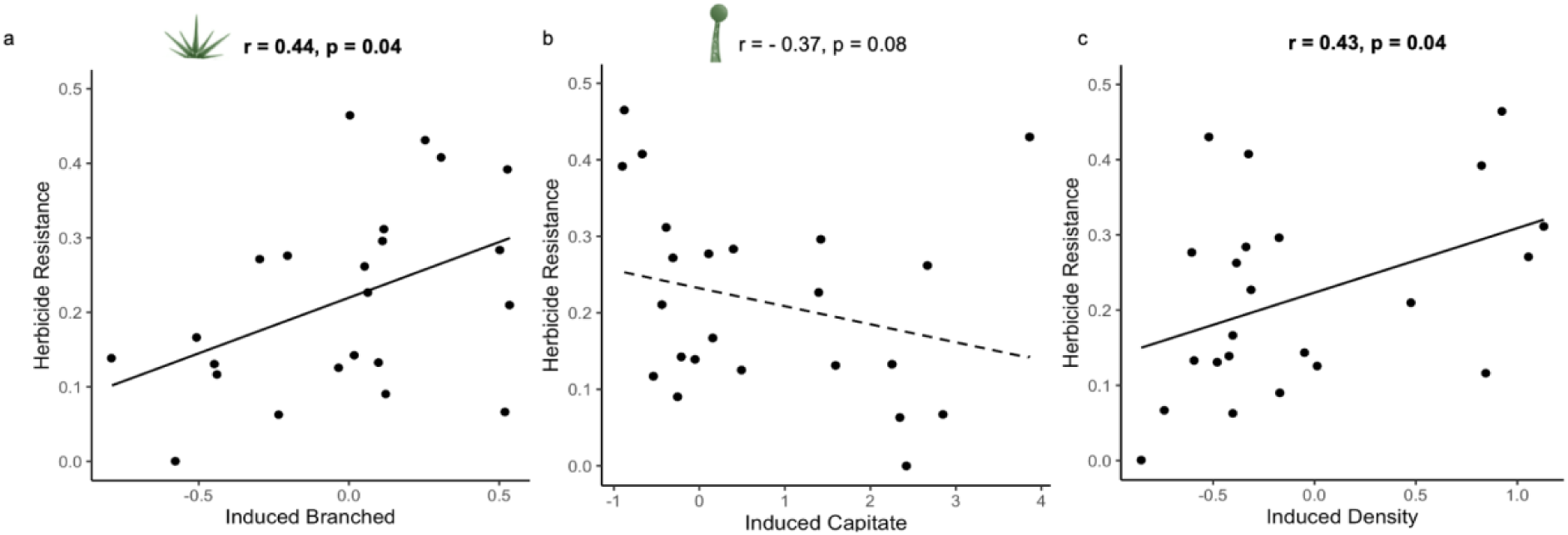
Relationships between herbicide resistance and (**a**) induced proportion branched and (**b**) induced proportion capitate and (**c**) induced total trichome density for *Abutilon theophrasti*. Shown are Pearson’s correlation coefficients; significant correlations are indicated in bold. Data points represent maternal line means. Induced trichome traits were measured as trichomes _damaged_ – trichome _control_, and herbicide resistance was measured as 1 – proportion of leaf yellowing.

**Figure 5.**
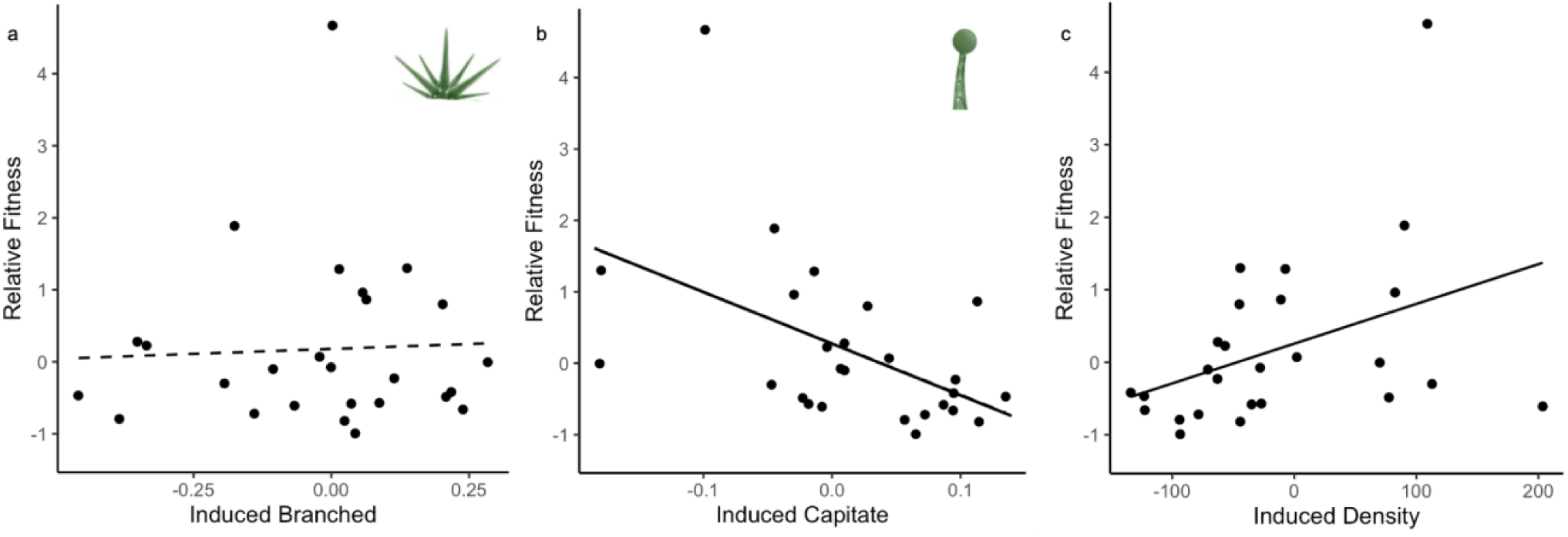
The relationship between relative fitness and (**a**) induced proportion branched (*S* = 0.10, p = 0.925) and (**b**) induced proportion capitate (*S* = -2.13, p = 0.045) and (**c**) induced total trichome density (*S* = 2.56, p = 0.018) in *Abutilon theophrasti*. Solid lines represent selection differentials in the presence of herbicide. Data points represent maternal line means; induced trichome traits were measured as damaged – control states, and herbicide resistance was measured as 1 – proportion of leaf yellowing.

There was no evidence of a relationship between the proportion of capitate trichomes and herbicide resistance (r = -0.37, p = 0.084, Fig 4B).

### Genotypic selection for induced trichomes

In the absence of herbicide, we found no evidence for selection acting upon any induced trait (branched: *S* = -0.03, p = 0.976; capitate: S = -0.39, p = 0.698, total density: S = -0.01, p = 0.99; STable 2). In the presence of herbicide, however, we found negative selection for induced capitate trichomes (*S* = -2.13, p = 0.045, Fig 4, STable 2) and positive selection for induced trichome density (*S* = 2.56, p = 0.018, Fig 4, STable 2). We found no evidence of selection on induced branched trichomes (*S* = 0.010, p = 0.925, Fig 4, STable 2) in the presence of herbicide.

In the absence of herbicide, we identified significant correlative selection acting branched trichomes (*γ* = 6.13, p = 0.006, STable 3), in those genotypes with the highest growth rate and a greater ability to alter the proportion of branched and capitate trichomes exhibited highest fitness. However, there was no evidence of correlative selection between growth rate and induced total density in this environment (*γ* = -0.87, p = 0.669, STable 3).

In contrast, in the presence of herbicide, there was evidence for correlative selection between growth rate and induced total density (*γ* = 5.71, p = 0.010, STable 3), in that genotypes with the highest fitness exhibited intermediate growth and high levels of induced total density (Fig 6.

**Figure 6.**
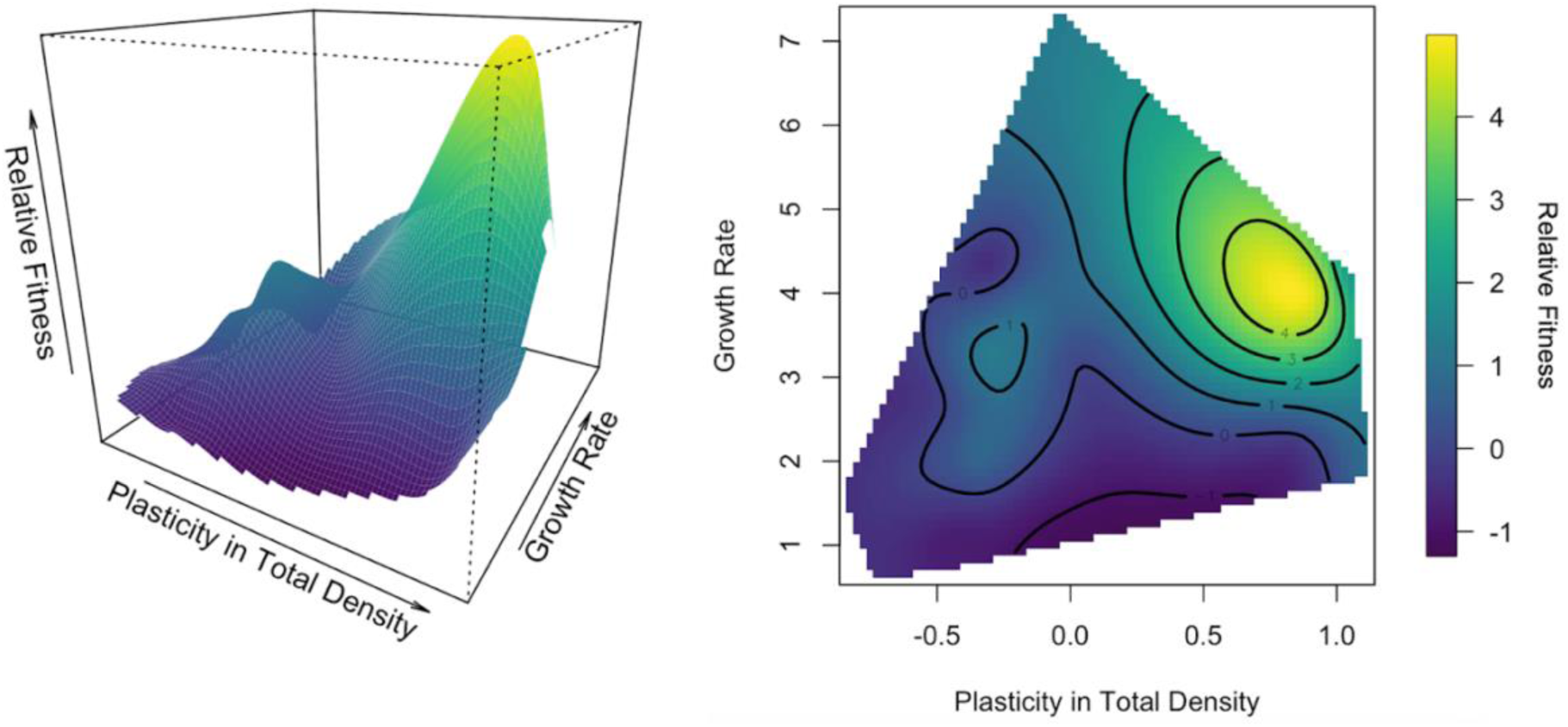
Fitness surface for correlative selection acting upon *Abutilon theophrasti* induced total trichome density and growth rate in the presence of herbicide (*γ* = 5.71, p = 0.010), tested using non-linear selection gradients (multivariate selection). Relative fitness is depicted by the color gradient; yellow is the highest fitness, blue is the lowest fitness, and green is intermediate.

There was no evidence of correlative selection acting upon growth rate and either induced branched (*γ* = -0.77, p = 0.070, STable 3) or induced capitate (*γ* = -2.58, p = 0.091, STable 3) trichomes in the presence of herbicide.

## Discussion

The goal of this study was to evaluate phenotypic plasticity in trichomes as a response to chemical stress from a common incrementally sprayed herbicide (glyphosate). Plasticity in defense traits such as trichomes may evolve in response to spatial and temporal variability in environmental stressors, allowing plants to balance the costs and benefits of defense, and are particularly favored when the presence of the stressor is unpredictable (Agrawal and Karban 1999; Karban and Baldwin 2007). We explored the cost and benefits of induced trichomes, which are apparent in the absence and presence of a damaging agent, respectively, reflecting a tradeoff between defense and reproduction (Rhoades 1979, Zangerl and Bazzaz 1992, Gavrilets and Scheiner, 1993).

In the broadest sense, plastic responses cause real-time modifications of resource allocation and can result in a range of defense strategies. For example, plants can induce phytochemicals (Baldwin and Schmelz 1994, Textor and Gershenzon 2009) and increase physical defenses such as thorns and trichomes (Young 1987, Milewski et al 1991, Gonzales et al 2008). Of the research on trichomes, ample work has examined environmental factors that induce trichome production to include: herbivory (Baur et al 1991, Agrawal 1999, Rautio et al 2002, Dalin and Bjorkman 2003), pathogens (Zhang et al 2020), water availability (Nagata et al 1999, Bosu and Wagner 2014, Vanhoutte et al 2017), UV radiation (Höglund and Larsson 2005), elevated CO_2_ (Karowe and Grubb 2011) and temperature (Ehleringer 1982, Perez-Estrada 2000, Shibuya et al 2016). However, few studies have identified the degree to which different trichome types respond plastically to novel stressors, acting as a multi-functionality stress-responsive trait (Sack and Buckley 2020). Here, we discuss our results conducted on velvetleaf (*Abutilon theophrasti*) as it relates to induced trichome production in response to herbicide exposure.

### Induced trichome production as a response to herbicide

Most plants have developed the ability to perceive and respond to external stimuli (Braam, 2005), and recent studies indicate that plant trichomes can directly sense external forces in anticipation of threats (Peiffer et al., 2009; Matsumura et al., 2022). Trichomes have been hypothesized to contribute to herbicide resistance (Devine et al., 1992; Baucom, 2019), but limited studies have empirically tested this idea. While we previously demonstrated that the proportion of branched trichomes is positively correlated with herbicide resistance in this species (Johnson and Baucom, 2024), suggesting a potential role as a constitutive resistance trait, the possibility that trichome plasticity also functions as an induced defense had not yet been explored.

In the present study, we show that trichomes are inducible in response to herbicide exposure, although the direction of the response varied. Across all plants, there was a general decrease in total trichome density on newly formed leaves following herbicide treatment, with no significant change in the proportion of trichome types. There were, however, significant maternal line effects in which half of maternal lines in this study had an increase in the proportion of capitate trichomes, our finding differs from previous work in M. guttatus which revealed that damage induced an increase in capitate trichome production. One important caveat is that we did not assess trichomes on the leaves that were directly exposed to the herbicide, which may exhibit different induction patterns. Despite an overall decrease in trichome density, however, we observed substantial variation among maternal lines. Specifically, 76% of maternal lines exhibited an increase in proportion of capitate trichomes, which aligns with previous work in *M. guttatus* which revealed that herbivory damage induced an increase in capitate trichome production (Holeski 2007). We also found that 24% of maternal lines exhibited an increase in total trichome density following herbicide exposure, and we detected a significant maternal line × treatment interaction for both traits. Moreover, those maternal lines with higher herbicide resistance showed stronger trichome induction overall—including increases in both total trichome density and the proportion of branched trichomes.

While we do not know the mechanistic basis of this positive correlation between herbicide resistance and induced trichomes, we hypothesize three potential reasons for this relationship. First, it is possible that genotypes that are more generally defended are likewise able to induce trichomes when damaged. In this scenario, which we imagine as agnostic to the particular damaging agent, the plant experiences stress that then triggers increased trichome production in genotypes that are better defended in a broad range of ways. In other words, individuals that are more herbicide-resistant also appear better able to mount a defense response, consistent with a generalizable inducible defense mechanism. Second, damage from an abiotic source, and specifically a chemical stressor like an herbicide, may trigger trichome production, potentially enhancing the plant’s ability to sequester chemicals within these structures, a process increasingly documented across diverse species (reviewed in Li et al 2022). Given that trichome vacuoles can occupy up to 95% of the total trichome volume (Gutiérrez-Alcalá et al., 2000), an increase in branched trichomes may be especially advantageous, as this type often possesses the highest vacuolar capacity (Calvert et al., 1985). Third, genes that influence trichome development in *A. thaliana* have known pleiotropic effects that impact other physiological functions (Bird and Gray 2003, Kirik et al 2004), thus it is possible that the induced trichomes we observed in velvetleaf could be linked to a response not measured in our experiment. While uncovering the exact mechanism underlying this relationship was outside of the scope of this study, further research is needed to determine if the induction in plant trichomes in more resistant genotypes is due to being more defended in general, having the ability to sequester chemicals for detoxification purposes, a pleiotropic relationship to an unknown response that we did not capture, or a combination of any of these explanations.

### Costs and benefits of trichome plasticity

The adaptive potential of damage-induced changes to trichome production are dependent on the benefits and costs of the phenotypic change (Dalin et al 2008). Positive correlations between induced increases in trichome traits and herbicide resistance suggests that plants would benefit from the induction of trichomes as long as the fitness benefits of this induction outweigh the costs of induction. Our linear selection analysis revealed that in the presence of herbicide there was positive selection acting upon the induced increase of total trichome density, but no evidence of selection acting upon the induced increase of proportion branched trichomes. This indicates that there are fitness benefits associated with the induced increase of total trichome density.

In the event of significant fitness cost, the pattern of selection would reveal fitness peaks in the absence of herbicide that correspond with the absence of the resistance character in question (*i.e*. induced total trichomes or induced increase of proportion branched trichomes) (Mauricio 1998). While many models assume that fitness costs increase linearly with allocation towards defense (Rhoades 1979, Simms and Rausher 1987), our results revealed no evidence of selection acting upon any of the induced trichome traits in the absence of herbicide, suggesting a lack of fitness costs. This finding aligns with several past studies in which costs for induced resistance had not been detected (Brown 1988, Simms 1992, Karban 1993, Mole 1994), but it also conflicts with other work that has identified costs of induced defenses (Tally et al 1999, Heil et al 2000, Cipollini 2002). Some explanations for the lack of costs may be that costs are too small, or that the costs may only be detectable in highly competitive environments (Simms 1992, Karban 1993). Here, we find that for induced trichome density, the net pattern of selection is positive, such that the benefits associated with trichome induction outweigh any cost.

### Other constraints on trichome plasticity

Plastic responses are expected to evolve when environmental cues provide a predictable indicator of appropriate adaptive phenotypes that exhibit genetic variation (Padilla and Adolf 1996; Reed et al. 2010). Thus, one constraint on the evolution of plastic defense responses would be the lack of genetic variation underlying such phenotypes. In this work we identified significant maternal line variation for induced total trichome density, as well as induced changes in the proportion of branched and capitate trichomes, indicating that these plastic trichome traits have the potential to evolve. While past work from other researchers has identified genetic variation in density, glandular, and non-glandular trichome types independently (Agrawal et al 2002, Holeski 2007, Ogran et al 2020), our study is one of few designed to detect genetic variation for induced trichome production for multiple trichome traits.

Past work tells us that in some cases induced plant defense can directly hamper reproductive output (Agrawal 1999, Redman 2001), and in other cases can directly hamper plant growth (Siemens 2002, Hermosa et al 2013). As such, we explored the possibility that allocation to induced defenses could come at a cost to plant growth. Following optimal defense theory, we predicted that in the presence of herbicide, selection should favor genotypes that allocated away from growth and toward induced defense, whereas in the absence of herbicide selection should favor genotypes that exhibited high plant growth and low levels of induced defense. We found different responses according to the type of trichome phenotype we considered. In the presence of herbicide, for total trichome density, we identified correlative selection favoring induced increases in the total number of trichomes and intermediate growth rates, indicating that induced trichomes come at the expense of increased growth. In contrast, in the absence of herbicide, we found no evidence of correlative selection acting upon induced total trichome density and growth rate, suggesting that plant growth is not limited in the absence of damage when trichome production is not induced.

In comparison, while we did not identify correlative selection acting on growth rate and either the proportion of branched and capitate trichomes in the presence of herbicide, in the absence of herbicide we found evidence of correlative selection in favor of both high growth rate and high inducibility for these trichome traits. This indicates that genotypes with the highest growth rate and high plastic ability to alter the proportion of trichome types have the highest fitness in the absence of herbicide. Overall, these results suggest that in the presence of herbicide, genotypes with intermediate growth and high plasticity in trichome production, but not plasticity in the proportion of trichome types, exhibit the highest fitness. These selection results align with past studies that indicate inducibility is negatively associated with species growth (Jacobsen 2022). Thus, while we did not find costs on induced trichomes in terms of fitness, the pattern of correlative selection in the presence of herbicide indicates a cost in terms of plant growth.

## Conclusion

Over the past four decades, much has been learned about the importance of trichomes in modulating leaf boundary dynamics, and recent developments suggest that we may be underestimating the eco-evolutionary significance of these plant appendages. As chemical stress is one of the main global drivers of environmental change, functional trait plasticity may play a key role in the persistence of plants in novel conditions. The current study examines induced trichome production as a reflection of the cost-benefit ratio associated with herbicide damage.

Our results provide evidence of a fitness benefit for induced trichome production in the presence of herbicide, demonstrating that induced responses can serve a stress responsive function against herbicide. It also provides evidence of evolutionary constraints in terms of correlative selection acting upon induced trichome production and reduced plant growth, indicating limiting factors that may impede, but not prevent, adaptive trichome plasticity. Future research should explore the multi-functionality of various trichome types and the potential for transgenerational trichome plasticity in ecosystems impacted by chemical stressors.

## Acknowledgments

The authors thank Michael Palmer and Jeremy Moghtader at the Matthaei Botanical Gardens for assistance with experiment logistics and herbicide applications. Many thanks to Kaira Liggett, Shantrell Tremmell, and Alanna Miyashiro who helped with field maintenance and with capturing pictures for herbivory estimates. Raj Gautam and Jordan Herman who helped collect and count seeds. We also thank Gregg Sobocinski at the University of Michigan Imaging Core for assistance with capturing trichome images.

## Author contributions

N.M.J and R.S.B. contributed to the experimental design, writing, and editing of this manuscript.

## Data Availability Statement

*The data underlying this article are available in* Github, at https://github.com/niajohnson1/HerbicideInducedTrichomes

## Supplementary Figures and Tables

**SFigure 1.**
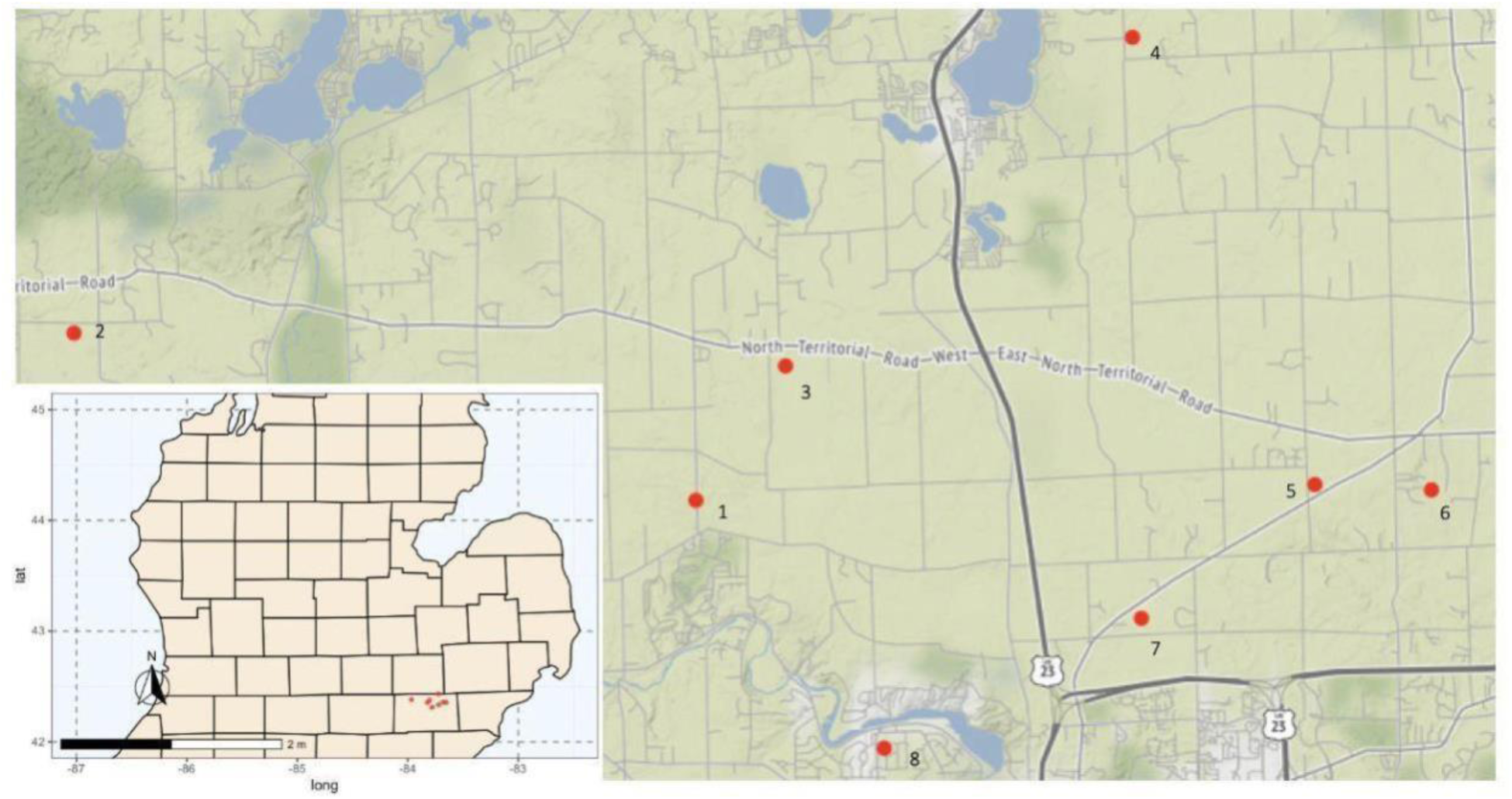
Locations of *Abutilon theophrasti* populations sampled and used for this study.

**STable 1.** Influence of herbicide treatment and maternal line on trichome traits (proportion branched, capitate, single, peltate, and total density) when observed in the field. Significant effects are indicated with asterisks: \**P* < 0.05, \*\**P* < 0.01, \*\*\**P* < 0.001.

| Trait | Treatment<br>(df=1) |  | Maternal Line<br>(df=1) |  | Maternal Line x<br>Treatment (df = 1) |  |
| --- | --- | --- | --- | --- | --- | --- |
| | F | p-value | $\chi^2$ | p-value | $\chi^2$ | p-value |
| Proportion Branched | 1.56 | 0.221 | 0.34 | 0.056 | 3.92* | 0.048* |
| Proportion Capitate | 1.94 | 0.168 | 0.00 | 1.000 | 5.64* | 0.018* |
| Proportion Single | 0.35 | 0.561 | 0.23 | 0.639 | 3.26 | 0.071 |
| Proportion Peltate | 0.44 | 0.512 | 0.08 | 0.767 | 0.71 | 0.399 |
| Total Density | 3.22 | 0.079 | 0.00 | 1.000 | 10.05** | 0.002** |

**STable 2.** Total selection (*S*) on *Abutilon theophrasti* induced trichome traits (proportion branched, proportion capitate, and total density) in the absence and presence of herbicide. Shown are selection differential values, standard errors, and p-values for traits in each treatment. Significant effects are indicated with asterisks: \**P* < 0.05, \*\**P* < 0.01, \*\*\**P* < 0.001.

| Phenotypic Selection (*S*) Differentials |  |  |  |  |  |  |
| --- | --- | --- | --- | --- | --- | --- |
| Trait | Control |  |  | Herbicide |  |  |
|  | *S* | SE | p-value | *S* | SE | p-value |
| Induced Branched | -0.22 | 0.15 | 0.826 | 0.10 | 0.47 | 0.925 |
| Induced Capitate | -0.38 | 0.04 | 0.707 | -2.13* | 0.11* | 0.045* |
| Total Induced Density | 0.50 | 0.09 | 0.625 | 2.56* | 0.37* | 0.018* |

**STable 3.**
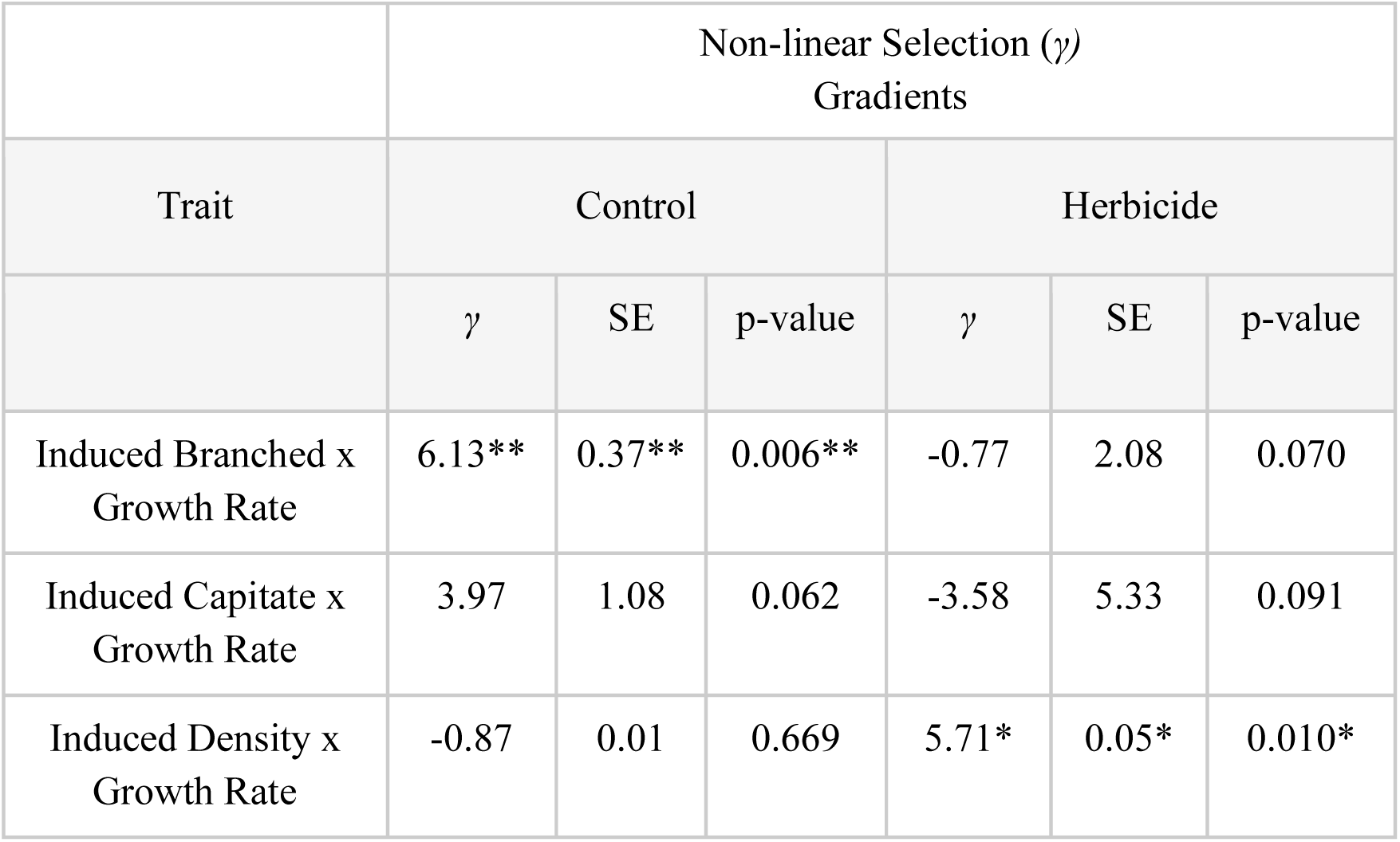
Correlative selection (*γ*) on *Abutilon theophrasti* induced trichome traits (proportion branched, proportion capitate, and total density) and growth rate in the absence and presence of herbicide. Shown are selection differential values, standard errors, and p-values for traits in each treatment. Significant effects are indicated with asterisks: \**P* < 0.05, \*\**P* < 0.01, \*\*\**P* < 0.001.

